# SCAMP - an open-source tool for profiling Alizarin Red-labelled calcification in fish larvae

**DOI:** 10.64898/2026.07.25.740697

**Authors:** Dávid Czimer, Panna Kaluzsa, József Kövendi, Luca Kamilla Li, Blanka Kapusi, Viola Pomozi, Krisztina Fülöp, Balázs Nagy, Csaba Benedek, András Váradi, Máté Varga

**Affiliations:** Department of Genetics, ELTE Eötvös Loránd University, Budapest, Hungary; Machine Perception Research Laboratory, HUN-REN Institute for Computer Science and Control, Budapest, Hungary; Institute of Molecular Life Sciences, HUN-REN Research Centre for Natural Sciences, Budapest, Hungary

**Author notes:** Correspondence (M.V.).

**Keywords:** zebrafish, ectopic calcification, notochord mineralization, image registration, Pseudoxanthoma elasticum

## Abstract

Quantifying skeletal mineralization phenotypes in larval fish is complicated by the natural curvature of the notochord and by sample-to-sample variability in orientation, staining and imaging. Consequently, many studies rely on summary measures such as vertebral counts or total stain intensity. Here we present SCAMP (Spinal Calcification & Mineralization Profiler), an open-source, GUI-based Python tool that computationally straightens the curved notochord of Alizarin Red S-stained fish larvae and generates standardized ARS-based mineralization profiles along the spinal axis. This approach reduces positional and shape variability, allowing direct quantitative comparison of Alizarin Red S fluorescence patterns within experimental cohorts processed and imaged under matched conditions, without requiring programming expertise. We validate SCAMP using a zebrafish model of Pseudoxanthoma elasticum (*abcc6a^elu15/elu15^*), recovering genotype-specific differences in the intensity, extent and spatial distribution of ectopic mineralization. Using SCAMP, we further show that inorganic pyrophosphate supplementation reduces ectopic intervertebral Alizarin Red S signal, alters the anterior-posterior distribution of mineralized regions in homozygous mutants, and increases signal at physiological chordacentrum positions. We also show that methylene blue, a routine antifungal additive in fish medium, is associated with reduced baseline Alizarin Red S-based mineralization endpoints, with the most pronounced effects observed in heterozygous controls. SCAMP is freely available and has the potential to be adapted to other fish species used in skeletal and mineralization research.

## Introduction

There are no two biological samples that are exactly the same. The stochasticity of biological processes results in constant variability on tissue, organ and organism level, therefore, even the most carefully collected specimens will differ in their minute details. The distortions that arise from methodological imprecisions during the analysis of the samples (i.e., differences in orientation, staining, mounting) introduce further imprecisions to the record of biological datasets. All these inaccuracies make the quantitative analysis of samples within and between experiments difficult.

Image standardization and registration, the spatial alignment of images into a common coordinate frame, offers a solution to this problem. The use of such standardized coordinate frames allows researchers to directly compare data acquired from different specimens, using different molecular labels and sometimes acquired even through differing modalities (e.g., light and electron microscopy) with one another. Such standardization can help to gather otherwise scattered, independent observations into comparable, quantitative datasets, making biological imaging reproducible, comparable and statistically poolable.

Over the past couple of years, several useful tools have been developed to aid such efforts. For example, brain imaging data in multiple species is routinely registered using the *ANTs* ecosystem (Tustison et al., 2021), histological data can be processed by *bUnwarpJ* (Arganda-Carreras et al., 2006) or *BigWarp* (Bogovic et al., 2016), and the *elastix* (Klein et al., 2010) toolbox and *VoxelMorph* (Balakrishnan et al., 2019) libraries are used to handle medical datasets.

Thanks to its small size, fecundity, tractable genetics and transparency, zebrafish (*Danio rerio*) has emerged over the past decades as an important model of biomedical research, and especially biomedical imaging (Lieschke and Currie, 2007; MacRae and Peterson, 2015; Patton et al., 2021; White et al., 2008). Indeed, many of the aforementioned image standardization and registration tools have been successfully applied to zebrafish research, mainly to create various reference brain atlases (Kunst et al., 2019; Marquart et al., 2017; Randlett et al., 2015; Ronneberger et al., 2012; Svara et al., 2022).

In larval teleosts, axial skeleton formation begins with the segmented mineralization of the extracellular notochord sheath (Grotmol et al., 2005; Inohaya et al., 2007). Specialized *entpd5*-positive notochord sheath cells, also termed chordoblasts, establish the metameric pattern of sheath mineralization along the anterior-posterior axis (Forero et al., 2018). The resulting mineralized rings are termed chordacentra and form in the absence of a cartilaginous anlage (Fleming et al., 2004). During subsequent development, chordacentra become surrounded by autocentra, which consist of bone produced by somite-derived scleroblasts. Accordingly, early notochord sheath mineralization should be distinguished from the later osteoblast-mediated formation of vertebral bone (Bensimon-Brito et al., 2012; Pogoda et al., 2018). The calcification of chordacentra is highly sensitive to the levels of systemic mineralization factors, and over the years several zebrafish models of human ectopic mineralization disorders have been identified and created by mutagenizing orthologous genes (Apschner et al., 2014; Czimer et al., 2021; Gils et al., 2018; Gils et al., 2022; Huitema et al., 2012; Mackay et al., 2015; Sun et al., 2020; Wynsberghe and Vanakker, 2023). The general feature of these models is precocious and often ectopic mineralization, that can be easily detected by Alizarin Red staining (Czimer et al., 2021; Gils et al., 2018; Gils et al., 2022; Mackay et al., 2015).

Due to the small size of the larvae and the ease of detection, these models and associated experimental paradigms have also been used to test the effect of potential drug candidates on the calcification process (Gils et al., 2022; Mackay et al., 2015; Sun et al., 2020). This is a particular strength of larval zebrafish models: mineralization phenotypes emerge early in development, well before they would be detectable in mammalian models, making larvae well suited to large-scale, rapid screening. However, the small size of the larvae that makes this possible also precludes many standard biochemical or metabolomic readouts, such as blood-based assays, that are routinely used to characterize mineralization disorders in mammals. As a result, precise and detailed quantification of the early visual (imaging-based) phenotype becomes disproportionately important. Yet current studies, rely on crude proxies for quantification: number of mineralized chordacentra or the overall intensity of Alizarin Red-stained areas are used to characterize the calcification process. This, partly, is due to the curved shape of the notochord and the slight stochasticity in the segmentation process. Although Alizarin Red S is widely used to visualize mineralized structures, its fluorescence intensity should be regarded as a relative imaging readout rather than a direct measure of absolute mineral content or mineral maturity.

Here, we present the Spinal Calcification & Mineralization Profiler (SCAMP), an open-source, GUI-based Python tool that computationally straightens the notochord in Alizarin Red S-stained fish larvae and generates standardized one-dimensional fluorescence intensity profiles along the spinal axis. Our approach relies on computationally straightening the naturally curved spinal axis prior to quantification, removing the positional and shape variability that otherwise confounds direct comparison of mineralization patterns between individual larvae.

We demonstrate the utility of SCAMP by examining the effects of inorganic pyrophosphate (PPi) supplementation on the extent and spatial distribution of ectopic calcification in *abcc6a* homozygous mutant zebrafish larvae. We also show that methylene blue, a standard disinfectant used in zebrafish media, can influence multiple quantitative features of spinal mineralization.

## Materials and methods

### Zebrafish husbandry and treatments

The *abcc6a^elu15^* mutant zebrafish were maintained in the animal facility of the Biology Institute of ELTE Eötvös Loránd University adhering to standard protocols (Aleström et al., 2019; Westerfield, 2000). Embryos and larvae used in our experiments were raised in E3 medium (5 mM NaCl, 0.17 mM KCl, 0.33 mM CaCl_2_ and 0.33 mM MgSO_4_ in distilled water) in a 28.5 °C incubator.

Embryos obtained from crosses between heterozygous *abcc6a^+/elu15^*and homozygous *abcc6a^elu15/elu15^* zebrafish were maintained in standard E3 embryo medium until 3 days post fertilization (dpf). At 3 dpf, larvae were individually transferred to 24-well plates containing 1 mL of treatment medium per well.

Four experimental treatment groups were established: untreated control (−MB/−PPi), pyrophosphate only (−MB/+PPi), methylene blue only (+MB/−PPi), and combined methylene blue and pyrophosphate treatment (+MB/+PPi). Treatment media were prepared using standard E3 medium supplemented with methylene blue (0.0001%, w/v) (E3 medium (for zebrafish embryos), 2011), disodium pyrophosphate hydrate (PPi) (1 mM), or both compounds, according to the assigned treatment group. All media were adjusted to pH 7.2– 7.5, prepared in 300 mL batches, aliquoted into sterile 30 mL conical tubes, and stored at −20°C until use. Before each medium change, aliquots were thawed and equilibrated to 28.5°C.

Larvae were exposed to their respective treatment media from 3 to 10 dpf. The treatment medium was completely replaced once daily by exchanging the entire 1 mL volume in each well with freshly thawed, pre-warmed treatment medium. Larvae were inspected daily during medium replacement for mortality and overt morphological abnormalities. These observations were used to identify gross treatment-associated toxicity but were not considered a formal quantitative developmental assessment. At 10 dpf, larvae from all treatment groups were collected simultaneously for fixation and subsequent Alizarin Red S staining.

Experiments were carried out in accordance with the Hungarian Act of Animal Care and Experimentation (1998, XXVIII) and with the directive 2010/63/EU of the European Parliament and of the Council of 22 September 2010 on the protection of animals used for scientific purposes. All protocols and experimental procedures used in this work were approved by the Hungarian National Feed Chain Safety Office (permit numbers PE/EA/2026-7/2017 and PE/EA/01615-2/2022) and the ELTE Eötvös Loránd University, Faculty of Science Animal Welfare Commission.

### Alizarin Red S staining and fluorescent image acquisition

At 10 dpf, larvae from all treatment groups were fixed simultaneously in 1 mL of 4% paraformaldehyde (PFA) in phosphate-buffered saline (PBS) at 4°C. Specimens were fixed overnight and, when immediate processing was not possible, maintained in the same fixative at 4°C for up to one week. Within each experiment, all treatment groups were kept in the fixative for the same duration and subsequently processed in parallel. Following fixation, each larva was washed twice in 1 mL distilled water for 20 min per wash. Specimens were then stained for 20 min at room temperature in 1 mL of freshly prepared 0.05% Alizarin Red S solution (VWR, CAS Number: 130-22-3). The working solution was prepared immediately before use by tenfold dilution of a 0.5% stock solution in distilled water. The stock solution was prepared in distilled water, adjusted to pH 7.0 with KOH, and stored at 4°C. Following staining, larvae were washed once in 1 mL distilled water for 20 min and once in 1 mL PBS for 20 min. All staining and washing steps were performed at room temperature with gentle agitation, using one larva per well. Specimens were protected from light throughout post-staining handling.

Stained larvae were subsequently embedded in 2% low-melting-point agarose and imaged on the same day using a Zeiss Apotome.2 structured illumination microscope (EC Plan-Neofluar 10×/0.30 objective, Axiocam 503 camera) with a fixed exposure time of 150 ms under identical acquisition settings for all samples. The use of the Zeiss Apotome.2 system was specific to the present study and is not a requirement of the SCAMP workflow. The image-processing and straightening procedures are not dependent on a particular microscope manufacturer or on structured-illumination imaging. Images acquired using other fluorescence microscope platforms can be analysed, provided that the spinal interval selected for analysis is captured with sufficient spatial resolution and quantitative dynamic range and that the z-step size is either retained in the image metadata or recorded separately.

Image acquisition for all specimens within an experiment was completed within 6 h after the end of the staining procedure. No stained specimens were stored overnight or imaged on a subsequent day. All samples were recorded using identical microscope settings, including objective lens, excitation and emission filter configuration, illumination intensity, camera exposure time and detector gain. Images were acquired in z-stack format to capture the complete three-dimensional distribution of mineralized structures within the spinal region. For the experiments reported in this study, raw image data were stored in Zeiss CZI format and used for all subsequent image-processing steps.

Following image acquisition, individual larvae were genotyped by allele-specific PCR as described before (Czimer et al., 2021) to distinguish wild-type, heterozygous, and homozygous *abcc6a*^elu15^ genotypes. Genotype information was subsequently used for data stratification and statistical analysis.

### ROI-Based Spatial Analysis

Quantitative image analysis was performed using ImageJ. All measurements were obtained from images processed using an identical, standardized thresholding procedure to ensure consistent segmentation of mineralized structures across all samples. The area and anterior– posterior (AP) position of each continuous mineralized region were subsequently extracted for statistical analysis. Statistical analyses were performed in R (Team, 2022). Multiple mineralized ROIs could be measured from the same larva, therefore, ROI-level observations were analysed using a linear mixed-effects model to account for within-fish clustering. ROI area was modelled as a function of genotype, scaled AP-axis position, and their interaction, with fish identity included as a random intercept (Area ∼ genotype × AP_axis_scaled + (1 | fishID)). AP-axis position was mean-centered and divided by its standard deviation before model fitting (Supplementary Table S1). The model was fitted by maximum likelihood using the *lme4* (Bates et al., 2015)and *lmerTest* (Kuznetsova et al., 2017) packages in R. Genotype-specific AP-axis slopes and their pairwise difference were estimated using *emmeans* (Lenth and Piaskowski, 2017). Model predictions with 95% confidence intervals were used for visualization. LOESS smooths were fitted directly to the ROI-level observations and were used exclusively for visualization. All statistical inference was based on the linear mixed-effects model. Scatter plots were generated using *ggplot2* (Wickham, 2016). Statistical significance was accepted at *p* < 0.05.

### SCAMP image processing pipeline

The complete workflow was implemented in Python (version 3.11) as a single application providing a graphical interface for image import, landmark placement, straightening and review. Image processing relied on *NumPy* (Harris et al., 2020) and *SciPy* (Virtanen et al., 2020) for numerical array operations, coordinate interpolation, and image resampling. Zeiss CZI files used in the present study were read using *AICSImageIO* (Brown et al., 2021), while TIFF-based image input and quantitative 16-bit TIFF output were handled using *tifffile* (Gohlke, 2022). *Pillow* (https://pillow.readthedocs.io) was used for general image handling, the graphical interface was built with *Tkinter*, and figures were rendered with *Matplotlib* (Hunter, 2007). Background regions of interest (ROIs) were defined either interactively within SCAMP or in ImageJ/Fiji and subsequently imported as ImageJ ROI files using the *roifile* library (Gohlke, 2022).

At minimum, input images must fully contain the spinal interval selected for quantitative analysis. In the present study, all SCAMP-derived fluorescence endpoints were calculated from the interval encompassing the seven most anterior chordacentra; therefore, imaging of the entire vertebral column was not required. The selected interval had to provide sufficient spatial resolution to distinguish chordacentrum and intervertebral signals, retain quantitative pixel intensities without substantial saturation or clipping, and include a signal-free region suitable for background estimation. The notochord boundaries also had to be sufficiently visible for reproducible landmark placement. For z-stack processing and depth normalization, the number of optical sections and the z-step size were obtained from the image data and associated metadata. When the z-step size was not available in the metadata, it could be supplied manually by the user.

The current implementation directly imports Zeiss CZI files as well as multidimensional TIFF and OME-TIFF image stacks. The downstream background correction, projection, straightening, and profile-extraction procedures are independent of the microscope manufacturer and do not require structured-illumination imaging or other specialized microscopy hardware. Images acquired using other fluorescence microscope platforms can therefore be analysed following lossless export to TIFF or OME-TIFF, provided that quantitative pixel values and stack organization are preserved and that the spatial information required for the selected processing workflow is retained in the metadata or supplied separately. Support for additional proprietary multidimensional image formats can be incorporated at the image-import layer without modification of the downstream quantitative analysis pipeline.

#### 1. Generation of background-corrected projection images

To obtain quantitative two-dimensional representations of mineralization-associated Alizarin Red S fluorescence, z-stack images were processed using a custom Python workflow. For each specimen, a manually defined background region of interest (ROI) was defined to estimate background fluorescence. SCAMP supports median-, mean-, and percentile-based estimation of background intensity. In the present study, the default median estimator was used. The median pixel intensity within the selected background ROI was calculated independently for each optical section and subtracted uniformly from all pixels in the corresponding section, thereby compensating for depth-dependent variation in background signal across the z-stack.

Following slice-wise background subtraction and clipping of negative values to zero, corrected optical sections were summed along the z-axis. The summed projection was divided by the total imaged depth, calculated as the number of optical sections multiplied by the z-step size in micrometres, yielding a background-corrected, Z-depth-normalized intensity projection. The resulting projection images were stored as quantitative 16-bit grayscale TIFF files and used for all subsequent analyses. A common quantitative intensity scale was retained among specimens acquired with identical microscope settings within the same experiment. For visual inspection only, an accompanying contrast-stretched copy of each image was generated. These preview images were not used in any quantitative analysis.

This approach minimized variability arising from background fluorescence and differences in stack depth while preserving quantitative intensity information.

#### 2. Definition of spinal reference geometry

The spinal region of larval zebrafish exhibits specimen-specific curvature. To define the spinal corridor for straightening, a series of manually positioned landmark triplets were placed along the rostro-caudal axis of each specimen. For all SCAMP-derived measurements, the analysed spinal interval was restricted to the seven most anterior chordacentra. Only the total number of mineralized chordacentra was assessed along the entire spinal region.

Each triplet consisted of a central point positioned approximately on the notochord midline and two auxiliary points placed beyond its dorsal and ventral margins to define the width of the analysed spinal corridor. The dorsal and ventral points were constrained to remain equidistant from the central point. Each triplet therefore defined a paired dorsal-to-ventral cross-section, including its local position, orientation, and width. The triplets were ordered according to the horizontal coordinate of their central points before image resampling.

Cubic-spline interpolation of the dorsal and ventral landmark sets was displayed in the graphical interface to guide landmark adjustment and was additionally used to define the source-corridor mask for quality control. These spline-interpolated coordinates did not contribute to the quantitative straightening transformation. Instead, the paired user-defined dorsal and ventral landmark coordinates were passed directly to the straightening procedure and linearly interpolated along centreline arc length to generate the dense resampling grid.

##### Geometry normalization by spinal straightening

To remove specimen-specific curvature, the guide-defined spinal corridor was resampled onto a straight rectangular coordinate grid. For each ordered landmark triplet, the centreline coordinate was calculated as the midpoint between the paired dorsal and ventral boundary points. The cumulative Euclidean distance between consecutive centreline coordinates was then used to parameterize position along the spinal corridor by centreline arc length.

The width of the straightened image was set from the total centreline arc length, with approximately one output column per source-image pixel unit along the centreline. The height of the output image was based on the median distance between the paired dorsal and ventral landmarks. Dorsal and ventral boundary coordinates were linearly interpolated as functions of centreline arc length. Each column of the target image represented a direct cross-section between the corresponding interpolated dorsal and ventral boundary coordinates.

For every target-grid position, the corresponding source-image coordinate was calculated by linear interpolation between the paired dorsal and ventral guide coordinates. Fluorescence intensities were sampled from the source image using first-order interpolation. No specimen-specific intensity normalization or thresholding was applied during straightening. The procedure therefore preserved the quantitative intensity scale of the input image, although individual pixel values and total intensity could change slightly because of geometric resampling. The resulting straightened fluorescence image was stored as a 16-bit TIFF for subsequent profile extraction.

#### 3. Characterization of larval mineralization profiles

Quantitative characterization of mineralization patterns was performed on straightened fluorescence images. For each image, fluorescence intensity values were integrated perpendicular to the longitudinal spinal axis by summing pixel intensities within each image column. This procedure converted the two-dimensional fluorescence distribution into a one-dimensional intensity profile representing the spatial distribution of mineralization-associated Alizarin Red S fluorescence along the analysed spinal interval.

For population-level visualization fluorescence intensity profiles were assembled into a common data matrix. Because the straightened images retained their specimen-specific arc-length-derived widths, the resulting one-dimensional profiles differed in length. For matrix export, shorter profiles were padded distally with missing values (NaN) to match the longest profile. No automatic intensity-based trimming was applied. Missing values were encoded as NaN rather than zero, ensuring that padded positions remained distinguishable from genuine zero-intensity measurements and were excluded from downstream quantitative analyses. Each specimen was associated with its corresponding experimental condition, which was stored together with the intensity profile in the exported data matrix.

The resulting matrix was exported in spreadsheet-compatible format and used to generate heatmap visualizations. Heatmaps provided a compact representation of fluorescence intensity distributions across entire experimental cohorts, enabling qualitative assessment of spatial variability and facilitating subsequent quantitative analyses. The combined matrix was written both as a comma-separated values (csv) file and as a spreadsheet (xlsx) workbook, each row labelled with the corresponding specimen and condition. When more than one experimental condition was present, samples were ordered by condition in the heatmap and annotated with a categorical colour strip and accompanying legend identifying each condition; with a single condition, an unannotated heatmap was produced. The heatmap was exported as a vector (pdf) and/or as a picture (png) file, with figure height scaled to the number of samples included.

### Experiment organization and data provenance

In SCAMP, all processing steps associated with a given experiment were organized under a unique experiment identifier of the form *yyyymmdd_<RANDOM suffix>*, which served as the name of the experiment root directory and was incorporated into every generated file name. Within this directory, workflow-specific subdirectories were automatically created to store background-corrected, Z-depth-normalized projection images (*background_subtracted*), straightened images (*straightened*), and downstream analysis outputs, including combined fluorescence intensity matrices and cohort heatmaps. Each output filename encoded the source specimen, the experiment identifier, and the assigned experimental condition, ensuring complete traceability of all intermediate and final data products throughout the workflow.

### Quantitative analysis and visualization of spinal mineralization in R

The fluorescence intensity matrices generated by SCAMP were imported into R (Team, 2022) for quantitative analysis and visualization. Data processing was performed using the *tidyverse* (Wickham et al., 2019) package, while statistical analyses were conducted using base R together with the *car* and *emmeans* packages where appropriate (Lenth and Piaskowski, 2017). Visualizations were generated using *ggplot2* (Wickham, 2016), and heatmaps were produced using the *ComplexHeatmap* package (Gu et al., 2016) with color scaling implemented through *circlize* (Gu et al., 2014). Prior to analysis, each one-dimensional fluorescence profile was resampled to 1000 equally spaced relative positions by linear interpolation. This procedure standardized the analysed spinal interval from 0% to 100% of its original profile length, enabling comparison of relative rostro-caudal signal distributions across specimens. Three metrics were calculated per profile. **Length-normalized integrated ARS fluorescence** was calculated as the sum of the 1000 interpolated profile values. Because all profiles contained the same number of normalized positions, this metric reflected average fluorescence intensity across the standardized spinal interval rather than absolute fluorescence integrated over the original physical profile length.

For the specimen-level analyses, a dataset-specific positive-fluorescence threshold was estimated from all finite, non-negative values of the length-normalized profiles. Values at or below the 60th percentile of the experiment-wide intensity distribution were treated as the operational low-intensity component. The threshold was defined as the median of this subset plus three scaled median absolute deviations (MAD; scaling constant = 1.4826). Profile positions with intensities strictly greater than this threshold were classified as positive. **Positive profile length** was calculated as the percentage of valid profile positions with intensities strictly greater than the threshold and was interpreted as the spatial extent of detectable above-threshold Alizarin Red S fluorescence. **Mean positive ARS fluorescence** was calculated as the mean intensity among above-threshold positions and represented the mean fluorescence intensity of the detected positive signal. Throughout this study, SCAMP-derived fluorescence metrics were treated as relative ARS-based mineralization readouts within each experiment and not as direct measures of absolute mineral content, composition, crystallinity or maturity. Thresholds were estimated separately for the MB/PPi experiment and the independent homozygous/heterozygous validation experiment because their fluorescence-intensity distributions and experimental cohorts differed. The primary thresholds were 611.05 a.u. for the MB/PPi experiment and 100.32 a.u. for the homozygous/heterozygous experiment.

The robustness of all threshold-dependent analyses was evaluated across a range of lower-tail quantile and MAD-based threshold specifications, with the 60% lower-tail quantile and three-MAD multiplier designated as the reference setting. For specimen-level endpoints, a single experiment-wide threshold was applied uniformly to all groups within each independently acquired experiment, and positive profile length and mean positive ARS fluorescence were recalculated under each alternative specification. For the ROI-like spatial-pattern analysis, treatment-specific thresholds calculated by pooling genotypes within each MB/PPi condition were used as the primary approach, while experiment-wide thresholds provided a more conservative sensitivity analysis. Robustness was assessed by comparing the direction and statistical interpretation of the principal model effects, treatment-specific genotype contrasts, AP-axis slopes, and PPi-associated slope shifts across threshold specifications. Because thresholds were estimated separately for independently acquired experiments, absolute threshold-dependent values were interpreted within, rather than directly compared between, experiments (Supplementary Tables S2A/1–4, S2B/1, S2C/1–5, and S2D/1–3).

For the primary intensity-based analysis of the MB/PPi experiment, each ARS-based mineralization endpoint was analysed using a three-factor linear model including genotype, MB exposure, PPi exposure, and all two– and three-way interactions (endpoint ∼ genotype × MB × PPi). Type III sums of squares were calculated using sum-to-zero contrasts. This factorial formulation allowed the effects of MB and PPi, and their potential dependence on genotype or co-treatment, to be evaluated separately. Genotype differences within each of the four MB/PPi treatment conditions were additionally assessed using two-sided Welch two-sample *t*-tests, with Benjamini–Hochberg correction across the four treatment-specific genotype contrasts for each endpoint. Treatment-versus-baseline contrasts within each genotype were also evaluated using two-sided Welch tests, with Benjamini–Hochberg correction across the five planned treatment contrasts for each endpoint and genotype (Supplementary Table S4).

To quantify treatment-associated changes in the spatial organization of mineralization, a complementary ROI-like pattern analysis was performed on the length-normalized fluorescence profiles. Positivity thresholds were estimated separately for each MB/PPi condition while pooling both genotypes within the respective treatment group. This treatment-specific thresholding was used exclusively to identify and compare continuous mineralized regions within each spatial profile and was not interpreted as a measurement of absolute fluorescence or absolute positive profile length. Continuous runs of above-threshold positions were extracted as individual positive regions. For each region, relative length was expressed as a percentage of the 1000-position normalized profile, and anterior–posterior position was defined by the centroid of the region on the normalized 0–100% spinal axis.

Relative region length was analysed using a linear mixed-effects model including genotype, MB, PPi, scaled anterior–posterior position, and all interactions as fixed effects, with fish identity included as a random intercept to account for the presence of multiple regions within individual larvae: region length ∼ genotype × MB × PPi × AP position + (1 | fish). Type III tests with Satterthwaite approximations were used for fixed-effect inference. Genotype– and treatment-specific AP-axis slopes were estimated from the fitted model, and PPi-associated changes in these slopes were tested within each genotype and MB condition using estimated marginal trends. Benjamini–Hochberg correction was applied across the four planned PPi slope contrasts (Supplementary Table S5).

For visualization, group mean fluorescence profiles were plotted as ribbon plots (95% CI shading) against normalized position, with the positivity threshold overlaid as a dashed line and shaded regions marking segments where the group mean exceeded it. To compare individual specimens, normalized profiles were assembled into a specimen × position matrix and hierarchically clustered by fluorescence pattern using *ComplexHeatmap* (Gu et al., 2016), with genotype annotations and a bar plot of length-normalized integrated ARS fluorescence per specimen displayed alongside. Spatial patterns of continuous threshold-positive regions were additionally visualized by plotting relative region length against anterior–posterior position using descriptive LOESS smooths with pointwise 95% confidence intervals. These curves were used for visualization only, while statistical inference was based on the linear mixed-effects model.

Because SCAMP fluorescence metrics are length-independent, the number of mineralized chordacentra was additionally scored manually as a complementary morphological measure of mineralization progression along the rostro-caudal axis, and analyzed separately from the SCAMP-derived metrics.

The scripts and related materials are available at our GitHub (https://github.com/danio-elte/SCAMP) and upon acceptance a timestamped version will be also uploaded to Zenodo.

## Results

Previously we have created and validated a novel *abcc6a* knock-out allele (*elu15*) to study Pseudoxanthoma elasticum (PXE) in zebrafish (Czimer et al., 2021). Similar to other reported *abcc6a* alleles (Gils et al., 2018; Mackay et al., 2015; Sun et al., 2020), homozygous *abcc6a^elu15/elu15^* mutants show elevated levels of calcification within the notochord, accompanied by ectopic intervertebral mineralization that can result in the fusion of adjacent chordacentra (Figure 1A).

**Figure 1:**
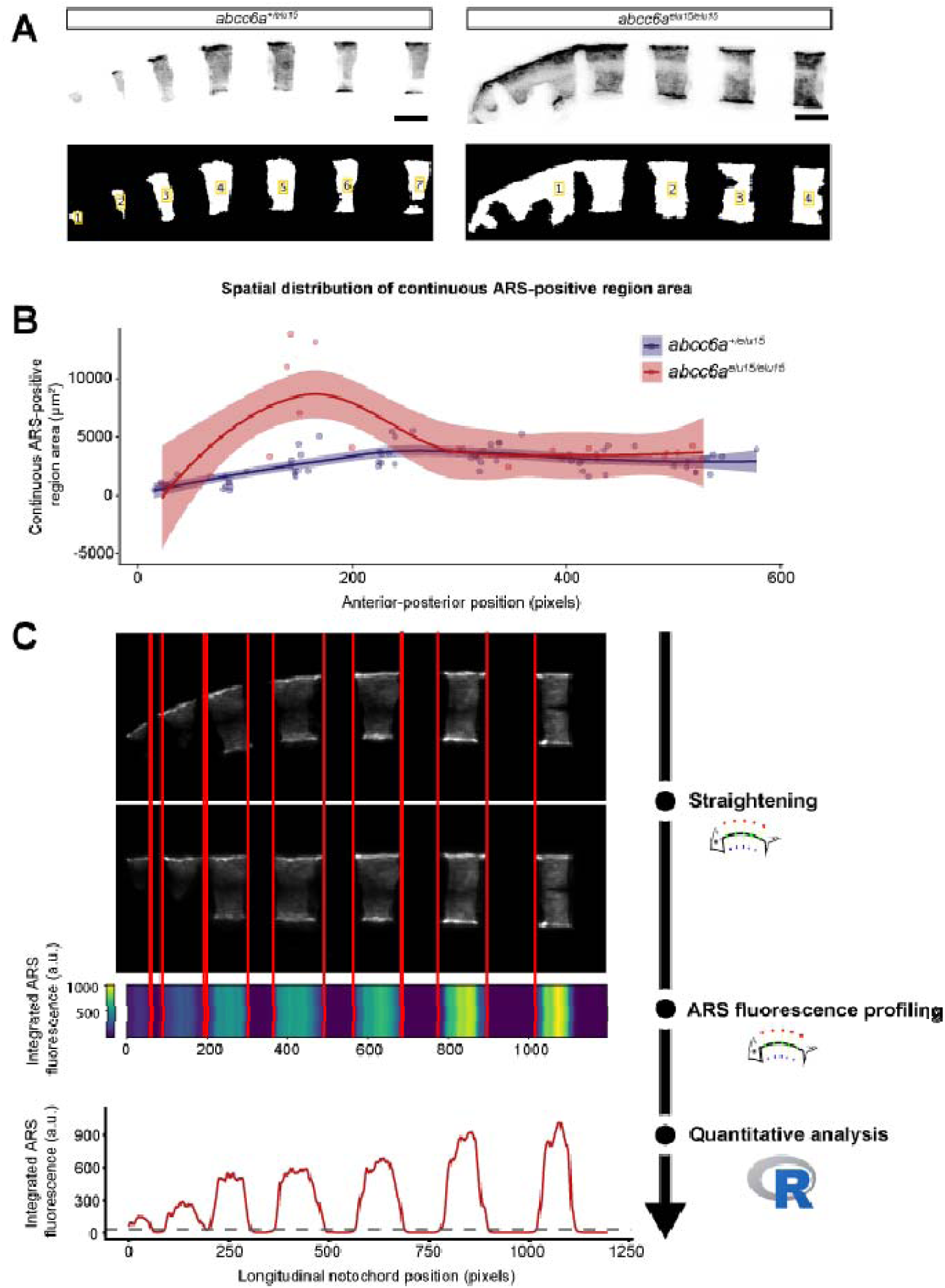
Limitations of region-of-interest-based analysis of ARS-positive regions in *abcc6a* mutants and overview of the SCAMP pipeline. (A) Representative Alizarin Red S (ARS) fluorescence images of the anterior notochord region in heterozygous *(abcc6a^+/elu15^*) and homozygous (*abcc6a^elu15/elu15^*) larvae. The corresponding thresholded binary masks are shown below, with numbered labels indicating the continuous ARS-positive regions identified as individual regions of interest (ROIs). In heterozygous larvae, spatially separated chordacentral ARS-positive regions can be identified as distinct ROIs. In homozygous mutants, ectopic intervertebral ARS fluorescence causes adjacent chordacentral regions to merge into larger continuous components, preventing reliable identification of individual chordacentra by connected-component-based ROI analysis. (B) Continuous ARS-positive region area plotted against anterior-posterior position in heterozygous and homozygous larvae. Lines represent descriptive LOESS smooths, and shaded ribbons indicate pointwise 95% confidence intervals. The LOESS curves are shown for visualization only; all statistical inference was based on a linear mixed-effects model with fish identity included as a random intercept. The relationship between continuous ARS-positive region area and anterior-posterior position differed significantly between genotypes (genotype × anterior-posterior position interaction: F1,81.06 = 24.63, P < 0.0001; 90 regions from 15 larvae). (C) Overview of the Python-based SCAMP workflow. The curved notochord image is computationally straightened, after which ARS fluorescence is integrated across the transverse dimension to generate a one-dimensional longitudinal fluorescence profile. Red guide lines indicate corresponding longitudinal positions across the processing steps. The resulting profile can subsequently be used for standardized quantitative and spatial analyses in R.

We initially quantified ectopic mineralization using an ROI-based analysis of Alizarin Red S-stained larvae (Figure 1B, Supplementary Table S1). The analysis included 90 continuous mineralized regions from 15 larvae. Because multiple regions could originate from the same larva, ROI areas were analysed using a linear mixed-effects model with fish identity included as a random intercept. The relationship between ROI area and AP-axis position differed significantly between heterozygous and homozygous mutant larvae (genotype × AP-axis interaction: F_1,81.06_=24.63, p<0.0001). These results demonstrate that the spatial distribution of continuous mineralized-region size along the AP axis differs between genotypes.

Although the ROI-based analysis detected genotype-dependent spatial differences, it represented each continuous mineralized region only by its area and position and therefore did not capture the complete fluorescence-intensity distribution along the curved notochord. We consequently developed a semi-automated workflow to computationally straighten the anterior notochord. The resulting linear geometry can be converted into a one-dimensional intensity profile, enabling direct comparison of the spatial distribution and intensity of mineralization-associated ARS fluorescence across specimens (Figure 1C).

The resulting SCAMP pipeline implements this workflow in four stages (Supplementary Video 1). First, fluorescence z-stacks of Alizarin Red S–stained larvae are imported, and a background region of interest is defined either interactively within SCAMP or by importing an ImageJ ROI file. The selected ROI is subsequently used for background estimation during intensity normalization. For each optical section, background intensity was estimated using the selected estimator (mean, median, or percentile-based). The estimated background value was subtracted from all pixels within the corresponding optical section, and negative intensities were clipped to zero. The background-corrected optical sections were subsequently summed along the Z axis and normalized by the total imaged Z-depth (number of optical sections × Z-step size, µm), yielding a background-corrected, Z-depth-normalized intensity projection. Second, the curved anterior notochord is annotated using a series of user-placed landmark triplets, each consisting of a midline point and two auxiliary points that define the dorsal and ventral boundaries of the spinal corridor. Third, a centreline is calculated from the paired dorsal and ventral boundaries and parameterized by cumulative arc length. The guide-defined spinal corridor is then resampled onto a straight rectangular grid, with each output column representing a dorsal-to-ventral cross-section at a defined centreline position. Pixel intensities were sampled from the source image using bilinear interpolation. No specimen-specific intensity rescaling or thresholding was applied during this transformation, although interpolation could modify individual pixel values. Finally, the straightened image is collapsed perpendicular to the long axis into a one-dimensional intensity profile, assembled across specimens into a condition-aware matrix that can be exported as a data table and/or can be visualized as a cohort heatmap. These outputs allow the user to compare the spatial distribution of mineralization-associated ARS fluorescence directly between individuals and genotypes.

First, we validated SCAMP using Alizarin Red S-stained heterozygous *abcc6a^+/elu15^* and homozygous *abcc6a^elu15/elu15^*larvae. The quantified profiles revealed genotype-dependent differences in the intensity and spatial pattern of ARS-labelled calcification (Figure 2A-C). By extracting further metrics from the dataset, we could confirm that length-normalized integrated ARS fluorescence and positive profile length were both greater in homozygous mutants than in heterozygous larvae (Figure 2D,E). Homozygous mutants showed a greater positive profile length than heterozygous larvae under the primary Q60_MAD3 threshold specification (Supplementary Table S3). This conclusion was robust across all nine tested threshold specifications: the estimated difference ranged from 20.16 to 26.94 percentage points and remained significant in every analysis (Welch p = 4.65 × 10 to 0.0179; Supplementary Table S2B/1). Interestingly, a trend toward greater inter-individual variability in length-normalized integrated ARS fluorescence was observed in homozygous mutants; however, this did not reach statistical significance (Brown–Forsythe Levene’s test, *P* = 0.0589; Supplementary Table S3).

**Figure 2:**
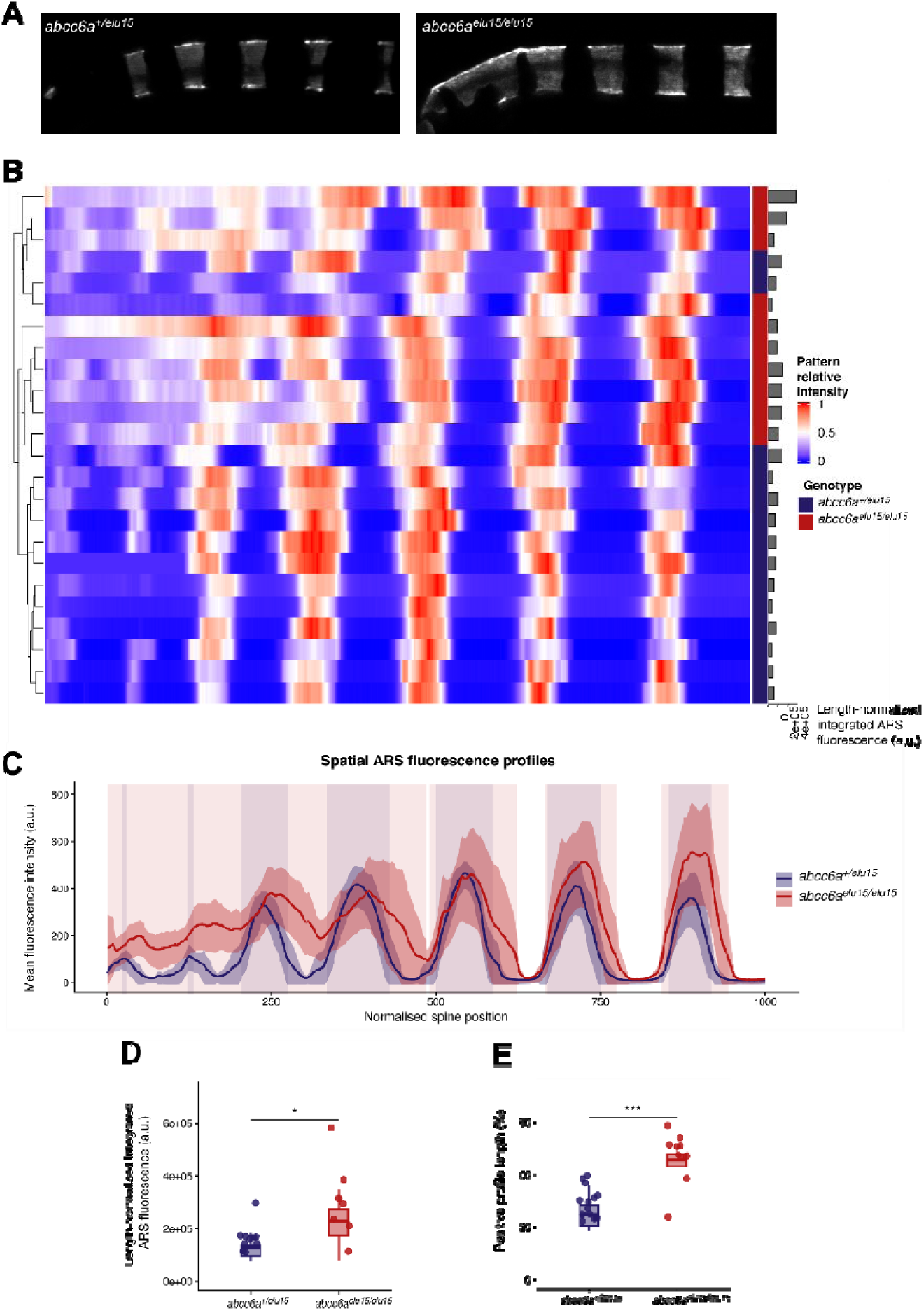
SCAMP enables the extraction of complementary spatial and quantitative Alizarin Red S fluorescence metrics from larval notochord profiles. (A) Representative maximum-intensity projections of mineralization-associated Alizarin Red S (ARS) fluorescence in the anterior notochord region of heterozygous (*abcc6a^+/elu15^*) and homozygous (*abcc6a^elu15/elu15^*) zebrafish. (B) Heatmap of spatial ARS fluorescence profiles generated by SCAMP. Each row represents one specimen, and profiles are hierarchically clustered according to their fluorescence distribution. Heatmap colours represent within-profile relative intensity scaled from 0 to 1 for visualization of spatial patterning; this scaling was not used to calculate the quantitative fluorescence endpoints. The annotation bar indicates genotype, and the adjacent bar plot shows the length-normalized integrated ARS fluorescence of each specimen. (C) Mean ARS fluorescence intensity profiles along the normalized spinal axis. Solid lines represent group means, and shaded ribbons indicate 95% confidence intervals. Blue and red represent abcc6a^+/elu15^ and abcc6a^elu15/elu15^ animals, respectively. The dashed horizontal line indicates the experiment-specific primary positivity threshold of 100.32 a.u., calculated as the median plus three scaled median absolute deviations (median + 3 × 1.4826 × MAD) using values at or below the 60th percentile. Shaded background regions mark spinal segments in which the group mean exceeded this reference threshold. These highlighted segments provide a descriptive group-level visualization and were not treated as independent inferential endpoints. (D) Length-normalized integrated ARS fluorescence. (E) ARS-positive profile length. The SCAMP-derived fluorescence endpoints were calculated from the interval encompassing the seven most anterior chordacentra. Data in panels D–E are shown as box-and-whisker plots overlaid with individual biological replicates. Boxes indicate the interquartile range (IQR), center lines represent medians, and whiskers extend to 1.5 × IQR. Statistical significance was assessed using two-sided Welch two-sample t-tests. Significance is indicated as ns (P ≥ 0.05), * (P < 0.05), ** (P < 0.01), *** (P < 0.001), and **** (P < 0.0001).

Next, we tested the utility of SCAMP for evaluating the effects of molecular treatments on spinal mineralization. Our previous studies showed that orally administered PPi can reduce ectopic calcification in mouse models (Dedinszki et al., 2017; Pomozi et al., 2017). We therefore examined whether supplementation of the larval medium with PPi similarly suppresses ectopic mineralization in *abcc6a^elu15/elu15^*fish larvae. Daily inspection during the treatment period revealed no overt morphological abnormalities or apparent treatment-associated increase in mortality in any of the MB/PPi treatment groups.

PPi supplementation visibly reduced ectopic intervertebral Alizarin Red S signal in homozygous mutant larvae and consequently reduced the genotype difference in positive profile length (Figure 3A,C). At the same time, PPi increased Alizarin Red S signal at physiological chordacentrum positions, particularly in heterozygous controls (Figure 3A,B). Consistent with this dual effect, the three-factor analysis identified significant PPi effects on length-normalized integrated ARS fluorescence, positive profile length, mean positive ARS fluorescence, and mineralized chordacentra count (Supplementary Table S4). A significant genotype × PPi interaction was detected for positive profile length (F = 23.74, *P* = 5.14 × 10), indicating that the effect of PPi on the spatial extent of mineralization differed between genotypes. PPi supplementation did not significantly alter the number of mineralized chordacentra in homozygous mutants. In heterozygous controls, the increase was most evident in the presence of MB, where PPi significantly increased the mineralized chordacentra count, further reducing the difference between genotypes (Figure 3D; Supplementary Table S4).

**Figure 3:**
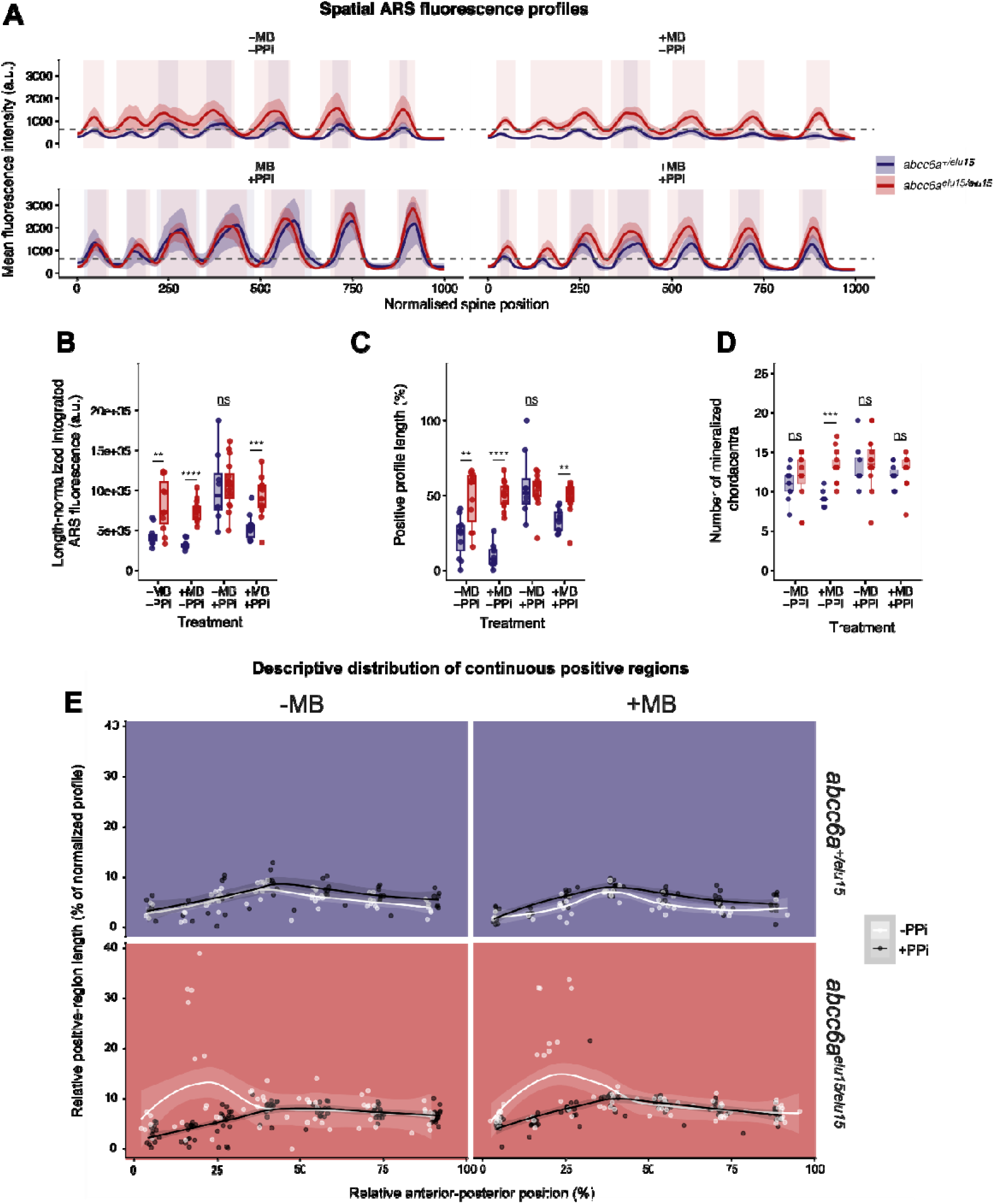
Inorganic pyrophosphate (PPi) and methylene blue (MB) alter the extent and spatial distribution of mineralization-associated Alizarin Red S fluorescence in zebrafish. (A) Mean Alizarin Red S (ARS) fluorescence intensity profiles along the normalized spinal axis for each treatment group. Solid lines represent group means, and shaded ribbons indicate 95% confidence intervals. Blue and red represent *abcc6a^+/elu15^* and *abcc6a^elu15/elu15^* animals, respectively. The dashed horizontal line indicates the experiment-specific primary positivity threshold of 611.05 a.u., calculated as the median plus three scaled median absolute deviations (median + 3 × 1.4826 × MAD) using values at or below the 60th percentile. Shaded background regions mark spinal segments in which the group mean exceeded this reference threshold. These highlighted segments provide a descriptive group-level visualization and were not treated as independent inferential endpoints. (B) Length normalized integrated ARS fluorescence. (C) ARS-positive profile length. (D) Number of mineralized chordacentra. The SCAMP-derived fluorescence endpoints in panels A–C were calculated from the interval encompassing the seven most anterior chordacentra, whereas chordacentrum counts were assessed separately and were not restricted to this interval. Data in panels B-D are shown as box-and whisker plots overlaid with individual biological replicates. Boxes indicate the interquartile range (IQR), center lines represent medians, and whiskers extend to 1.5 × IQR. The effects of genotype, MB, PPi, and their interactions were assessed using three-factor linear models with Type III analysis of variance. Genotype contrasts within individual MB/PPi treatment conditions were evaluated using two-sided Welch two-sample t-tests with Benjamini-Hochberg correction across the four treatment specific comparisons for each endpoint. Statistical significance is indicated as ns (P ≥ 0.05), * (P < 0.05), ** (P < 0.01), *** (P < 0.001), and **** (P < 0.0001). (E) Relative lengths of continuous threshold-positive regions plotted against their anterior-posterior positions. Each point represents one continuous threshold-positive region, and multiple regions could originate from the same animal. Lines represent descriptive LOESS smooths, and shaded ribbons indicate pointwise 95% confidence intervals. Panels are separated by genotype and MB exposure; white and black lines represent the absence and presence of PPi, respectively. The LOESS curves are shown for visualization only; all statistical inference was based on the linear mixed-effects model reported in Supplementary Table S5, in which animal identity was included as a random intercept.

The ROI-like spatial-pattern analysis further showed that PPi markedly altered the anterior– posterior distribution of continuous mineralized regions in homozygous mutants (Supplementary Table S5). Without PPi supplementation, larger mineralized regions were concentrated toward the anterior part of the analysed spinal interval, consistent with the prominent anterior ectopic intervertebral mineralization visible in untreated mutants. PPi supplementation eliminated or reversed this anterior bias, shifting the distribution of larger mineralized regions toward more posterior positions. This PPi-associated shift was significant both in the presence and absence of MB, whereas no corresponding change was detected in heterozygous controls (Supplementary Table S5).

Together with the fluorescence profiles and representative images, these findings indicate that PPi reduces the characteristic anterior ectopic mineralization pattern of homozygous mutants while preserving or increasing the ARS-based mineralization readout at physiological chordacentrum positions.

The opposing spatial effects of PPi may reflect differences between pathological intervertebral mineralization and physiological mineralization at chordacentrum sites. One possible explanation for the opposing effects of PPi is that non-hydrolysed PPi remains available to inhibit ectopic mineral deposition in normally non-mineralizing regions of the notochord, while PPi hydrolysis in the medium or within the larvae increases Pi availability. Developing chordacentra may preferentially use this additional Pi because they provide a physiological environment that supports mineralization, with local phosphatase activity potentially contributing to this process. Together, these effects could explain the reduction in ectopic intervertebral mineralization alongside the increased ARS-based mineralization readout at physiological chordacentrum positions and the posterior shift observed after PPi treatment. However, PPi hydrolysis, Pi availability, and phosphatase activity were not measured directly, and this interpretation remains to be tested.

We additionally examined the effect of methylene blue (MB) on spinal mineralization. MB is commonly used as an antifungal disinfectant in zebrafish embryo medium (Aleström et al., 2019; Westerfield, 2000), but recent evidence indicates that it can also influence mitochondrial function and metabolism in developing larvae (Nipu et al., 2025). Because mitochondrial dysfunction has been implicated in PXE (Lofaro et al., 2020; Nollet and Vanakker, 2022), we investigated whether MB influences the mineralization phenotype of our zebrafish PXE model or modifies the observed response to PPi supplementation.

At the routinely used concentration of 3.1 µM, MB exposure was associated with lower spinal Alizarin Red S-based mineralization endpoints, predominantly in heterozygous controls. These larvae showed lower length-normalized integrated ARS fluorescence, positive profile length, mean positive ARS fluorescence, and number of mineralized chordacentra following MB exposure, whereas the corresponding changes in homozygous mutants were smaller and less consistent across endpoints (Figure 3; Supplementary Figure S1; Supplementary Table S4). This difference between genotypes was most evident for positive profile length: MB markedly reduced the positive profile length representing the spatial extent of detectable ARS fluorescence in heterozygous controls but had little effect in homozygous mutants, resulting in a significant genotype × MB interaction (F, = 9.00, *P* = 0.00355). Thus, the association between MB exposure and the ARS-based mineralization endpoints was endpoint– and genotype-dependent, with the clearest response observed in heterozygous controls.

The magnitude of the MB-associated changes also varied between PPi conditions, although evidence for a general modifying effect of PPi was limited. An MB × PPi interaction was detected only for length-normalized integrated ARS fluorescence, and no genotype × MB × PPi interaction was found for any endpoint (Supplementary Table S4). Therefore, PPi co-treatment did not consistently alter the response to MB across genotypes and mineralization measures.

These findings raise the possibility that MB influences the observed mineralization phenotype through its reported effects on cellular redox state or mitochondrial metabolism, both of which have been implicated in PXE pathophysiology (Nollet and Vanakker, 2022). The pronounced mineralization phenotype caused by loss of Abcc6a function may make the more modest and endpoint-specific effects of MB less apparent in homozygous mutants. However, oxidative stress and mitochondrial activity were not measured directly in the present study.

Sensitivity analyses using experiment-wide thresholds ranging from 398.11 to 948.28 a.u. showed that the principal conclusions for positive profile length and mean positive ARS fluorescence were robust across all nine threshold specifications. The significance classifications of all terms in the three-factor genotype × MB × PPi models remained unchanged, as did the significance classifications of all treatment-specific genotype contrasts. For positive profile length in the −MB/+PPi condition, the small, non-significant genotype difference changed direction under two threshold specifications but remained non-significant throughout. Thus, none of the biological conclusions depended on the selected positivity threshold (Supplementary Tables S2A/1 and S2C/1–5). Because the genotype × MB and genotype × PPi interactions for positive profile length remained significant under every threshold specification, genotype differences in this endpoint were interpreted in the context of the individual MB/PPi conditions (Supplementary Tables S2C/2–3). The principal conclusions of the ROI-like spatial-pattern analysis were similarly robust. PPi produced a significant shift in the AP-axis distribution of mineralized regions in homozygous mutants under all treatment-specific threshold specifications, whereas no corresponding shift was detected in heterozygous controls (Supplementary Table S2D/2).

## Conclusions

To aid the quantification and analysis of phenotypes that affect the mineralization process in the notochord, we have created a user-friendly open-source tool, Spinal Calcification & Mineralization Profiler (SCAMP). The tool provides an easy-to-use graphical interface, that makes adoption easy, as no formal programming skills are required.

Alizarin Red S fluorescence provides a relative imaging readout of mineralization under standardized experimental conditions but does not map linearly onto absolute mineral content or mineral maturity. Accordingly, SCAMP-derived intensity measurements were interpreted as comparative ARS-based mineralization endpoints within experiments, rather than as direct measurements of mineral mass, composition or crystallinity.

We tested SCAMP with our recently developed PXE model, the *abcc6a^elu15/elu15^* mutant line, where loss of Abcc6a function leads to precocious calcification of the chordacentra and results in ectopic calcification loci (Czimer et al., 2021). Both the initial ROI-based analysis (Figure 1B) and the subsequent SCAMP profiles revealed a pronounced genotype-dependent spatial pattern, with homozygous mutants displaying larger continuous mineralized regions toward the anterior portion of the analysed notochord (Figure 2).

As expected, PPi supplementation reduced the ectopic mineralization phenotype, and the principal threshold-dependent conclusions were robust to alternative positive-threshold specifications (Figure 3; Supplementary Tables S2C/1–5 and S4). The ROI-like spatial-pattern analysis further showed that PPi eliminated or reversed the anterior enrichment of large continuous mineralized regions characteristic of homozygous mutants. This PPi-associated shift was detected both in the presence and absence of MB and remained significant across all primary treatment-specific threshold scenarios (Supplementary Tables S2D/2 and S5). At the same time, PPi enhanced the mineralization-associated ARS fluorescence at physiological chordacentrum sites. One possible explanation is that non-hydrolysed PPi inhibits ectopic mineral deposition in normally non-mineralizing regions of the notochord, whereas PPi hydrolysis in the medium or within the larvae increases Pi availability for mineralization at developing chordacentra. Local phosphatase activity may also contribute to this process; however, this mechanism was not directly tested here. Several observations argue against a generalized developmental delay as the principal explanation for the PPi-associated changes. The number of mineralized chordacentra showed only modest and condition-dependent differences among treatment groups, while the fluorescence-based effects were genotype– and position-dependent. Most notably, PPi reduced ectopic intervertebral Alizarin Red S signal in homozygous mutants while preserving or increasing signal at physiological chordacentrum positions. Such spatially opposing effects are not readily explained by a uniform delay in development.

We also identified an unexpected association between MB exposure and several Alizarin Red S-based mineralization endpoints. At the routinely used concentration, MB exposure was associated with reductions in several ARS-based mineralization endpoints, with the most consistent effects observed in heterozygous controls. This finding indicates that routine medium additives can influence quantitative mineralization phenotypes and should therefore be carefully standardized in pharmacological and genetic experiments. MB may affect mineralization through its reported effects on cellular redox state or mitochondrial metabolism (Nipu et al., 2025), however, these mechanisms were not directly examined in the present study. No overt morphological abnormalities or apparent increase in mortality were observed following MB exposure. Moreover, the MB-associated changes were endpoint– and genotype-dependent rather than uniform across the experimental groups. Nevertheless, because developmental progression was not assessed using an independent quantitative staging endpoint, a subtle effect of MB on developmental timing cannot be excluded completely.

Because methylene blue has antimicrobial activity, we considered whether its presence might indirectly influence PPi availability by limiting microbial degradation in the treatment medium. In an ancillary stability experiment, however, PPi concentrations remained close to the initial 1 mM concentration throughout the seven-day observation period even in the absence of methylene blue (Supplementary Figure S2). Although a modest MB-dependent difference in the relative temporal slopes was detected, neither MB condition showed a significant fish-versus-medium-control slope difference after correction for multiple comparisons. Most importantly for the interpretation of the treatment experiments, no substantial depletion of PPi was observed in fish-conditioned medium without methylene blue. These observations support that PPi remained sufficiently stable over the treatment period even in the absence of methylene blue, although the experiment was not designed as a formal equivalence or non-inferiority test.

These two observations underscore the advantages of standardized image processing for monitoring ectopic and endogenous calcification in the notochord of fish larvae.

Finally, we also want to point out that our tool is not restricted to zebrafish, as it can be also used on the larvae of other fish species, where either genetic mineralization models can be created – such as medaka (*Oryzias latipes*) (Lleras-Forero et al., 2020; To et al., 2011) – or aquaculture conditions might result in known developmental skeletal defects, such as Atlantic salmon (*Salmo salar*) (Witten et al., 2016) and Senegalese sole (*Solea senegalensis*) (Azevedo et al., 2017).

## Data availability

Raw image data created for this study has been deposited in the BioImage Archive (https://www.ebi.ac.uk/biostudies/bioimages/studies/S-BIAD3836). Scripts related to the SCAMP pipeline can be found on our GitHub repository (https://github.com/danio-elte/SCAMP) and a timestamped version will be also deposited in Zenodo upon the acceptance of the manuscript.

## Author contributions

Conceptualization: CB, AV, DC, MV.

Methodology: DC, KF, VP, BN, CB, AV, MV.

Investigation: DC, PK, LKL, BK.

Software: JK, DC, MV.

Resources: VP, KF, AV, MV.

Writing – original and revised text: DC, MV.

Visualization: DC, MV.

Funding acquisition: AV, MV.

Supervision: MV.

## Acknowledgements

We would like to thank Anita Rácz for fish care, and members of the ELTE Fishgenetics Group for valuable comments.

