## Supplementary Figures for "SCAMP - an open-source tool for profiling Alizarin Red-labelled calcification in fish larvae"

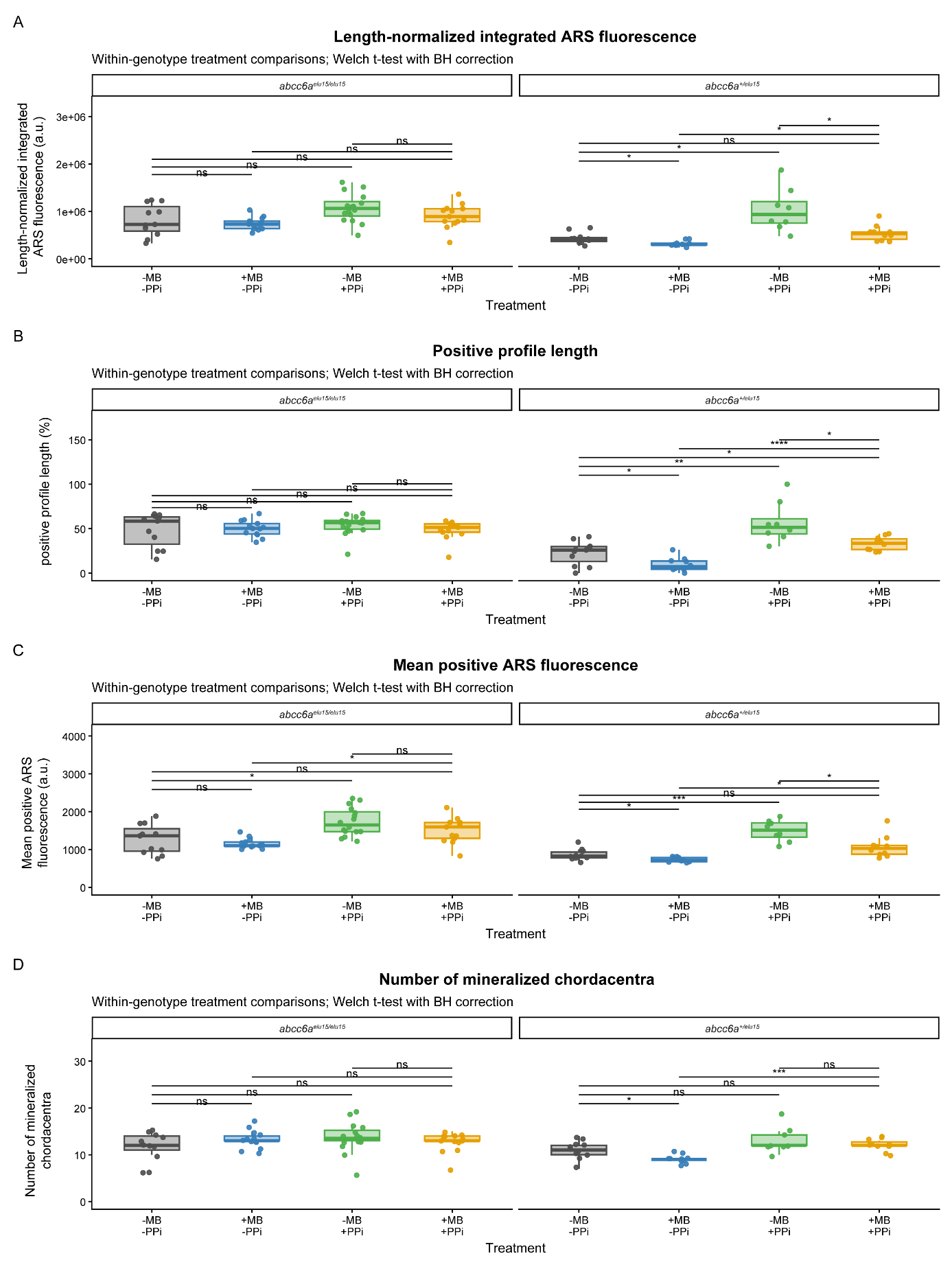

**Supplementary Figure S1: Within-genotype effects of methylene blue and inorganic pyrophosphate on ARS-based spinal mineralization endpoints.** (A) Length-normalized integrated ARS fluorescence. (B) ARS-positive profile length. (C) Mean positive ARS fluorescence. (D) Number of mineralized chordacentra. Heterozygous (abcc6a+/elu15) and homozygous mutant (abcc6aelu15/elu15) larvae are shown separately across the four methylene blue (MB) and inorganic pyrophosphate (PPi) treatment conditions. The experiment-specific positivity threshold used for the endpoints in panels B and C was 611.05 a.u. For panels A–C, fluorescence-based endpoints were calculated from the spinal interval encompassing the seven most anterior chordacentra, whereas chordacentrum counts were assessed separately and were not restricted to this interval. Data are shown as box-and-whisker plots overlaid with individual biological replicates. Boxes indicate the interquartile range (IQR), center lines represent medians, and whiskers extend to 1.5 × IQR. Five planned treatment contrasts within each genotype were evaluated using two-sided Welch two-sample t-tests, with Benjamini–Hochberg correction applied across the five comparisons separately for each endpoint and genotype (Supplementary Table S4). Statistical significance is indicated as ns (P ≥ 0.05), * (P < 0.05), ** (P < 0.01), *** (P < 0.001), and **** (P < 0.0001).

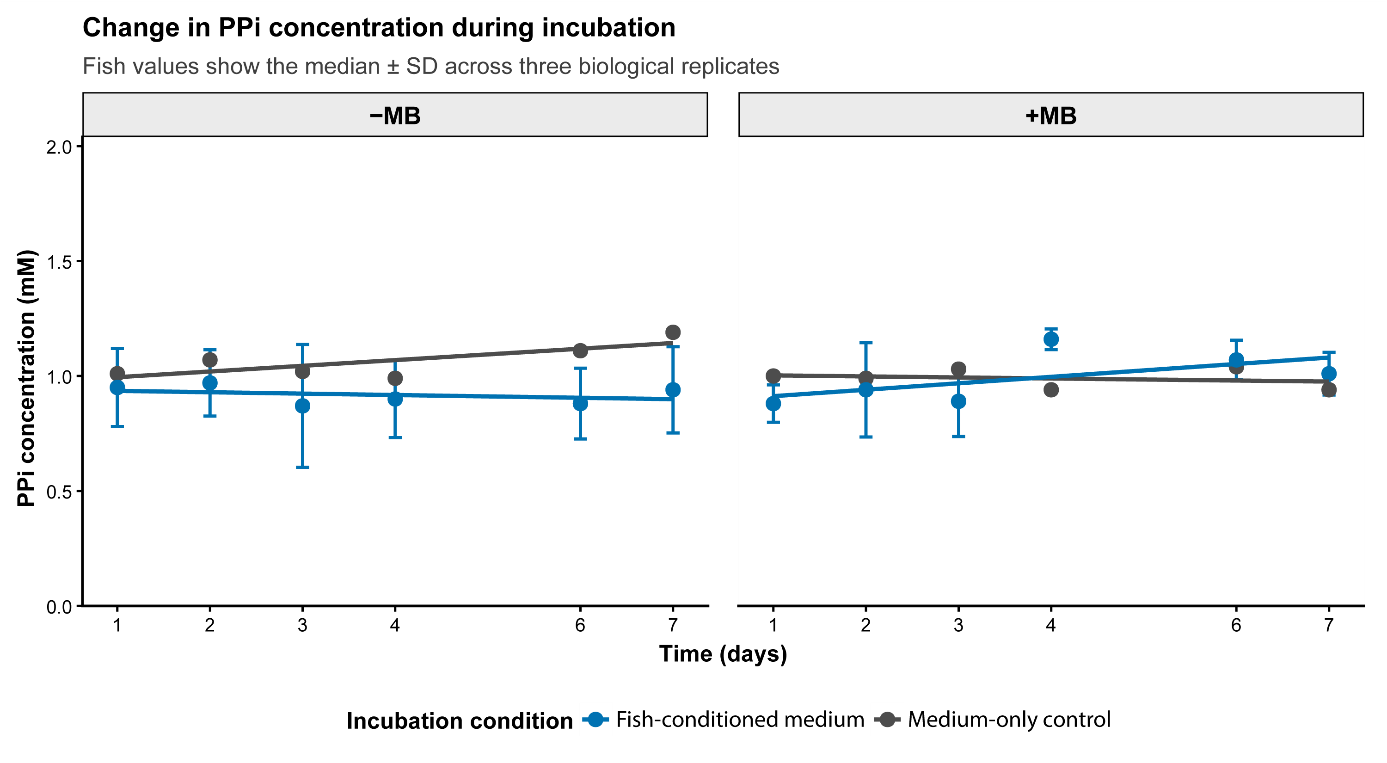

**Supplementary Figure S2: Stability of PPi in the treatment medium in the presence and absence of methylene blue.** PPi concentration was monitored for seven days in medium-only controls and in media incubated with fish, either without (−MB) or with (+MB) methylene blue. Fish-conditioned values are shown as the median ± SD across three biological replicates after technical measurements were summarized within each fish. Medium-only controls represent the median of technical measurements and are shown without error bars. Lines indicate descriptive linear trends and are included for visualization only. Statistical inference was based on a linear mixed-effects model including incubation time, the presence of fish and MB treatment. A modest MB-dependent difference in the fish-versus-control temporal slopes was detected (time × fish exposure × MB interaction: estimate = 0.066 relative PPi units per day, 95% CI: 0.003–0.129, P = 0.040), although the fish-versus-control slope difference was not significant within either MB condition after multiple-testing correction. PPi concentrations remained close to the initial 1 mM concentration throughout the observation period, including in fish-conditioned medium without methylene blue, supporting sufficient PPi stability under the applied treatment conditions.
